# Status and targets for rebuilding the three major fish stocks in Lake Victoria

**DOI:** 10.1101/2020.10.26.354639

**Authors:** Laban Musinguzi, Mark Olokotum, Vianny Natugonza

## Abstract

We determined fisheries management reference points for three major fish stocks in Lake Victoria (Nile tilapia, Nile perch and Dagaa) for Uganda and the whole lake. The aim was to ascertain stock status and define reasonable objectives and targets for rebuilding to sustainable levels. Dagaa was found to be healthy in Uganda and the whole lake but tending to overfished status. In Uganda, the stock status of Nile tilapia and Nile perch was recruitment impaired but tending more towards collapsed and overfished status respectively. In the whole lake, the stock status of Nile tilapia and Nile perch was collapsed and overfished respectively with the latter tending more towards recruitment impaired. Estimates of maximum sustainable yield (MSY) showed that catches could be increased under good management. Rebuilding the Nile tilapia and Nile perch stock biomasses to MSY level (B_msy_) could respectively increase the catches above the current level by 9.2% and 29.5% in Uganda and by 72.8% and 15.1% in the whole lake. The immediate objective for fisheries management should be to rebuild biomass for the Nile tilapia and Nile perch stocks to B_msy_. Elimination of illegal fishing practices has proved to be effective. In addition, management needs to keep catches at low levels until biomass for the stocks is ≥B_msy_ for at least three consecutive years.

## Introduction

Uganda and Tanzania have since 2017 strengthened enforcement of fisheries regulations to end illegal fishing and improve stocks of Lake Victoria. With ~1 million tons of fish produced annually, Lake Victoria supports the world’s biggest inland fishery useful for foreign exchange, employment and direct sustenance of >4 million people (Marshall & Mkumbo 2011). These benefits have for a long time been threatened by high fishing pressure (Njiru et al, 2007; Nyamweya et al. 2020), justifying the strengthened enforcement.

The countries strengthened the enforcement by deploying their respective defense forces, Fish Protection Unit (FPU) in Uganda and the multisector task force in Tanzania (Mudliar, 2018; NPA, 2019). These partially or fully replaced previous institutional arrangements such as beach management units which were considered ineffective because of corruption (Nunan et al. 2018). In Uganda, the enforcement demonstrated determination to end illegal fishing practices as the deployment was followed by a total stop on all forms of illegalities. The ineffective institutional arrangements were replaced, Illegal gears and crafts destroyed, fishers forced out of near shore areas.

Positive outcomes have been observed from the enforcement. Data from fishery independent surveys conducted since 2017 show that the biomass of Nile perch (*Lates nilotics* (Linnaeus, 1758)), the most important commercial fish species in the lake has improved and was at its record high since 2010 (Hydro-acoustics Regional Working Group, 2019). Interestingly, 48% of the increase was recorded between 2018 and 2019, with the largest increase recorded in the Ugandan part followed by Tanzania. The increase in biomass was least in Kenya where enforcement was not strengthened. The surveys further showed that although Nile perch was still dominated by individuals under the size at which recruitment to the fishery occurs, there was a record increase in the proportion of fish at the preferred size. These observations suggest that good management in Lake Victoria can pay off.

With all due respect, the enforcement is ongoing with no consideration of fisheries management reference points, lacking clear management objectives and targets beyond the elimination of the illegal fishing gears and practices. To contribute to effective enforcement, we estimated fisheries management reference points for major commercial fish species to act as a basis of adopting evidence-based fisheries management objectives and targets.

The reference points determined for the whole lake and the Ugandan part of the lake clarify on the status of the stocks before the commencement of the enforcement. They are not only useful for substantiating objectives and targets for the enforcement but are also indispensable for evaluating its effectiveness. At the global scale, this assessment is commensurate with calls to increase assessment of inland fish stocks to support responsible inland fisheries (Cooke et al. 2016, FAO & MSU, 2015; FAO, 2020).

## Methods

### Approach and stocks assessed

The reference points were based on two methods, a Monte Carlo method (CMSY) and a Bayesian state-space implementation of the Schaefer production model (BSM). A brief background to these methods is provided here while more details are available in Froese et al. (2017). The methods are built on a principal that catch from a species is produced by its biomass and productivity such that if two of the three parameters are known, production models can be used to estimate the other. The CMSY uses catch and productivity to estimate biomass. The method uses prior ranges of productivity and current biomass (B) relative to unexploited biomass (k) (B/k) at the start, intermediate and end of a time series to detect productivity and unexploited biomass pairs with corresponding biomass estimates that are compatible with observed catches. The BSM on the other hand uses catch and biomass data to estimate productivity. The methods are integrated with other empirical formulae to estimate the reference points including maximum sustainable yield (MSY), fishing mortality rate F at MSY (F_msy_), biomass required to support MSY (B_msy_), relative stock size (B/B_msy_) and exploitation (F/F_msy_).

We estimated reference points for three major stocks in Lake Victoria at two spatial scales: the whole lake and the Ugandan part of the lake. Nile perch, *Oreochromis niloticus* (Linnaeus, 1758) (Nile tilapia) and *Rastrineobola argentea* (Pellegrin, 1904) locally known as Dagaa are the three major fish species supporting commercial fisheries in Lake Victoria. The species are responsible for >88.7% (estimated from catch used in this study) of catches from Lake Victoria. Dagaa is the most important by weight, followed by Nile perch and Nile tilapia. Nile perch supports fish processing industries that export to foreign markets including the European Union and is the most important by value.

### Data requirements, sources and application to CMSY

To estimate the reference points, abundance and catch data were required at the two spatial scales. For the whole lake, we estimated the reference points using two indices of abundance i.e. absolute biomass and fishery independent catch per unit effort (CPUE). This provided an opportunity to evaluate the usefulness of both sets of data which are available for Lake Victoria. The reference points for the Ugandan part of the lake were based on CPUE only.

The absolute biomass was obtained from Nyamweya et al. (2016) who simulated the biomass based on catches and hydrodynamics of the lake using ecosystem models. This was available to 2015, starting from 1965, 1968 and 1971 for Nile perch, Dagaa and Nile tilapia respectively. The CPUE was from hydroacoustic (Nile perch and Dagaa) and trawl (Nile tilapia) surveys and was only consistent for the species since 1999. The CPUE was restricted to 2015 beyond which no catch data are available.

Catch data used for the whole lake was partly available from Nyamweya et al. (2016) and was supplemented with data from the archives of the National Fisheries Resources Research Institute. The archives were the sources of the catch data for the Ugandan part.

Productivity of a stock is reflected in CMSY as prior ranges of intrinsic rate of population increase (r) which are derived by classifying resilience of species available in FishBase into r values (Froese & Pauly, 2015; Froese & Pauly, 2019). The resilience of Dagaa is high and that of Nile tilapia and Nile perch is medium. Their respective r ranges are 0.6-1.5, 0.2-0.8 and 0.2-0.8 (Froese & Pauly, 2019; Froese et al. 2017).

The B/k prior ranges depend on depletion status of stocks: very strong depletion (0.01 – 0.2), strong depletion (0.01 – 0.4), medium depletion (0.2 – 0.6), low depletion (0.4 – 0.8), and nearly unexploited (0.75 – 1.0) (Froese et al. 2019). These are required for the start, intermediate and end year of the time series. We harnessed trends in biomass and catches over the time series to set the B/k ranges for the species (Figure 1).

**Figure 1.**
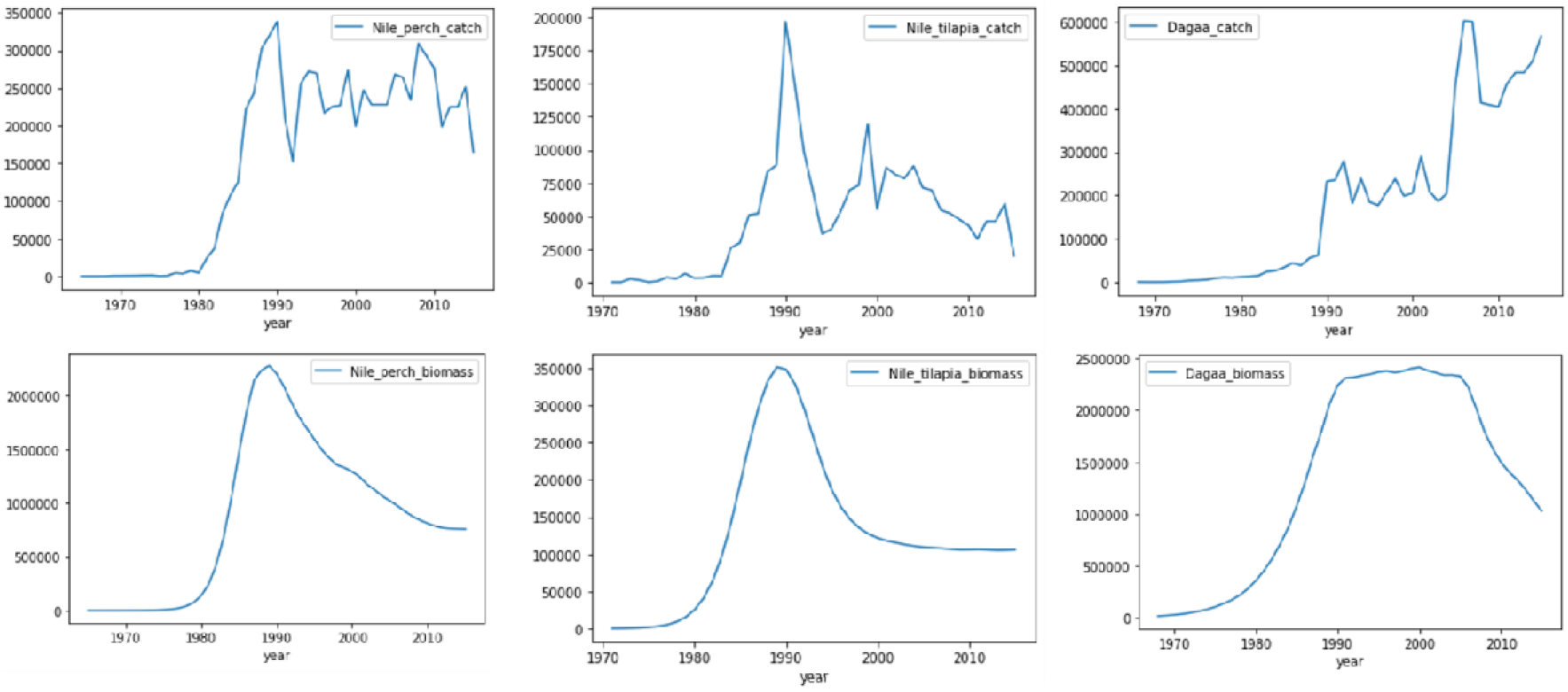
Trends in catches and absolute biomass for the three major commercial fish species on Lake Victoria. Catches are for the whole lake. Absolute biomass from Nyamweya et al. (2016).

The start years for the Nile perch, Nile tilapia and Dagaa were 1965, 1971, 1968 respectively in the first scenario for the whole lake using absolute biomass, and 1999 for the second whole lake scenario using CPUE and the Ugandan part. The end year was 2015 in both scenarios.

For the first whole lake scenario, the B/K ranges at the start years were set at low depletion for all the stocks (Table 1). In these years, the stocks were at the initiation fishery development phase and the low depletion enabled the future development of their fisheries (Hilborn & Walters, 1992; Figure 1). At the end, the Nile perch fishery had reached the decline phase of fishery development (Hilborn & Walters, 1992; Figure 1) which was characterized by high fishing effort (Nyamweya et al. 2020). For this reason, we set the B/k priors to strong depletion. For Nile tilapia, by 2015, although the catch in the whole lake was not the lowest ever, it was 10.3% of the historical maximum, guiding us to set the B/K prior to very strong depletion. For Dagaa, the catch was increasing albeit decreasing biomass. Given its high turnover rate and high fishing pressure (Mangeni-Sande et al. 2019), we set its B/K priors to medium.

**Table 1.**
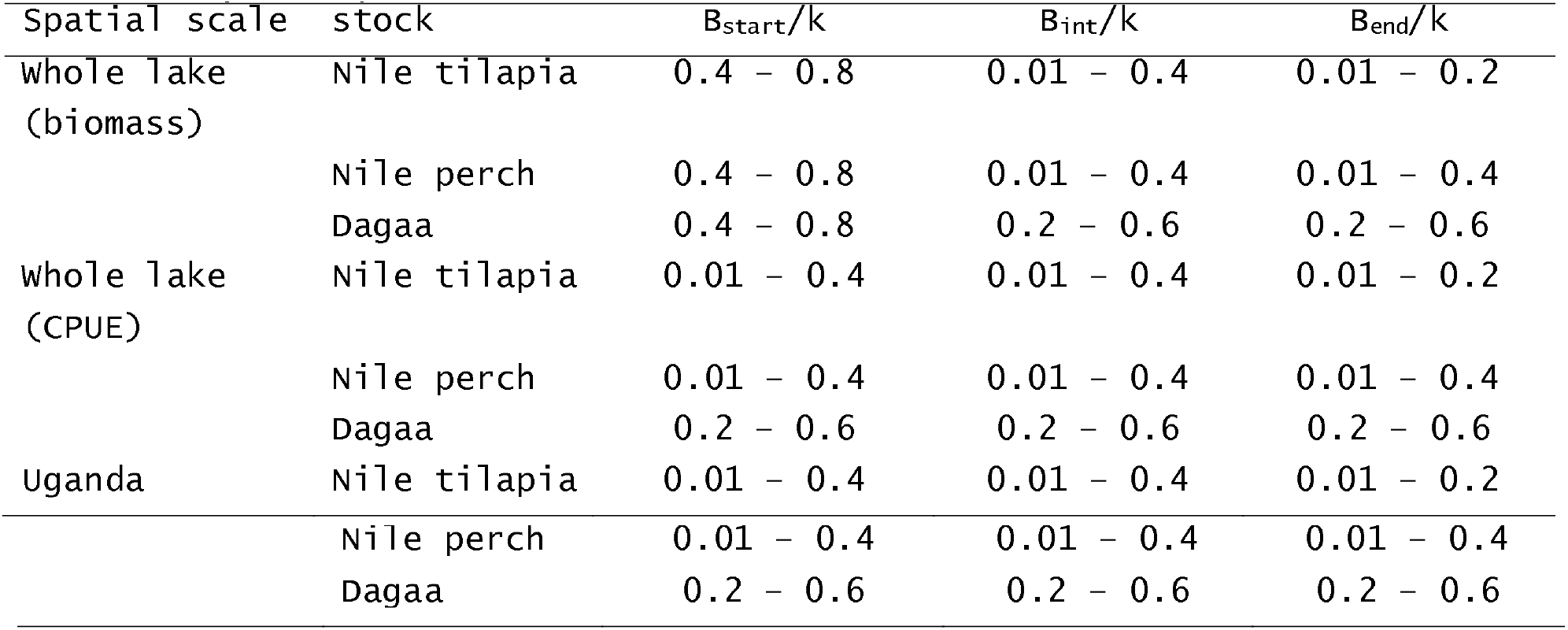
Relative stock biomass (B/K) prior ranges used for the stocks. The ranges depend on the depletion status of stocks: very strong depletion (0.01 – 0.2), strong depletion (0.01 – 0.4), medium depletion (0.2 – 0.6), low depletion (0.4 – 0.8), and nearly unexoloited CO.75 – 1.0) (Froese et al. 2019)

| Spatial scale | stock | B<sub>start</sub>/K | B<sub>int</sub>/K | B<sub>end</sub>/K |
| --- | --- | --- | --- | --- |
| whole lake<br>(biomass) | Nile tilapia | 0.4 – 0.8 | 0.01 – 0.4 | 0.01 – 0.2 |
|  | Nile perch | 0.4 – 0.8 | 0.01 – 0.4 | 0.01 – 0.4 |
|  | Dagaa | 0.4 – 0.8 | 0.2 – 0.6 | 0.2 – 0.6 |
| whole lake<br>(CPUE) | Nile tilapia | 0.01 – 0.4 | 0.01 – 0.4 | 0.01 – 0.2 |
|  | Nile perch | 0.01 – 0.4 | 0.01 – 0.4 | 0.01 – 0.4 |
|  | Dagaa | 0.2 – 0.6 | 0.2 – 0.6 | 0.2 – 0.6 |
| Uganda | Nile tilapia | 0.01 – 0.4 | 0.01 – 0.4 | 0.01 – 0.2 |

|  |  |  |  |
| --- | --- | --- | --- |
| Nile perch | 0.01 – 0.4 | 0.01 – 0.4 | 0.01 – 0.4 |
| Dagaa | 0.2 – 0.6 | 0.2 – 0.6 | 0.2 – 0.6 |

The intermediate year is a year in the development of a fish stock such as when biomass, exploitation or recruitment was low or high (Froese et al. 2019). We set the intermediate years for the stocks at years when biomass was highest i.e. 1989 for Nile perch, 1991 for Nile tilapia and 2000 for Dagaa. For the Nile perch and Nile tilapia stocks, the intermediate years were at the time fishery development was declining (Figure 1), prompting us to set the B/k priors to strong depletion (Table 1. For Dagaa, medium depletion was selected.

In the scenarios of CPUE as the index of abundance, we set the B/k priors at the start (1999) for the whole lake and Ugandan part at strong depletion for Nile perch and Nile tilapia, and medium depletion for Dagaa based on guidance from trends in biomass. The end year B/K priors were maintained as above for both spatial scales. The intermediate years for the Ugandan part were set at 2005 for all the stocks, corresponding to a period when fishing effort, catch, and CPUE were highest or lowest. The corresponding intermediate B/k priors were set at strong depletion for Nile perch and Nile tilapia, and medium depletion for Dagaa. This was similar for the whole lake only that the intermediate years were 2005 for Nile tilapia, 2008 for Nile perch and 2007 for Dagaa.

Finally, a recent period of at least 5 years when catch and abundance were relatively stable or had similar trends is required for determining catchability coefficient to relate CPUE to biomass. The default of the last 5 years was selected for all the stocks except Dagaa in the whole lake whose catches and abundance had different trends (Figure 1). We chose a period from 2000-2004 when biomass and catches for Dagaa in the whole lake were stable.

The CMSY/BSY were implemented in R using the code for the methods (Froese et al. 2019). Data used are available online (Musinguzi, 2020). Palomares et al. (2018) established an approach of classifying fish stocks as collapsed, recruitment impaired, overfished, or healthy, basing on estimates of B/B_msy_. This was used to define the status of the stocks assessed.

## Results and discussion

For the whole lake, the two indices of abundance used returned comparable estimates of the fisheries reference points because values in both scenarios were largely overlapping and falling within each other’s confidence intervals (Table 2; supplementary table 1). For this reason estimates with CPUE as the index of abundance were adopted for further consideration. In addition, CPUE is the most preferred for the methods used (Froese et al. 2017) and its results were more precautionary for most of the stocks. Tables 2 and 3 present the estimates of the reference points and stock status determined for the whole lake and Ugandan part respectively. Figures 2, 3 and 4 illustrate the reference points and status for Nile perch in Uganda as an example, with illustrations of other stocks available in supplementary material 1.

**Figure 2.**
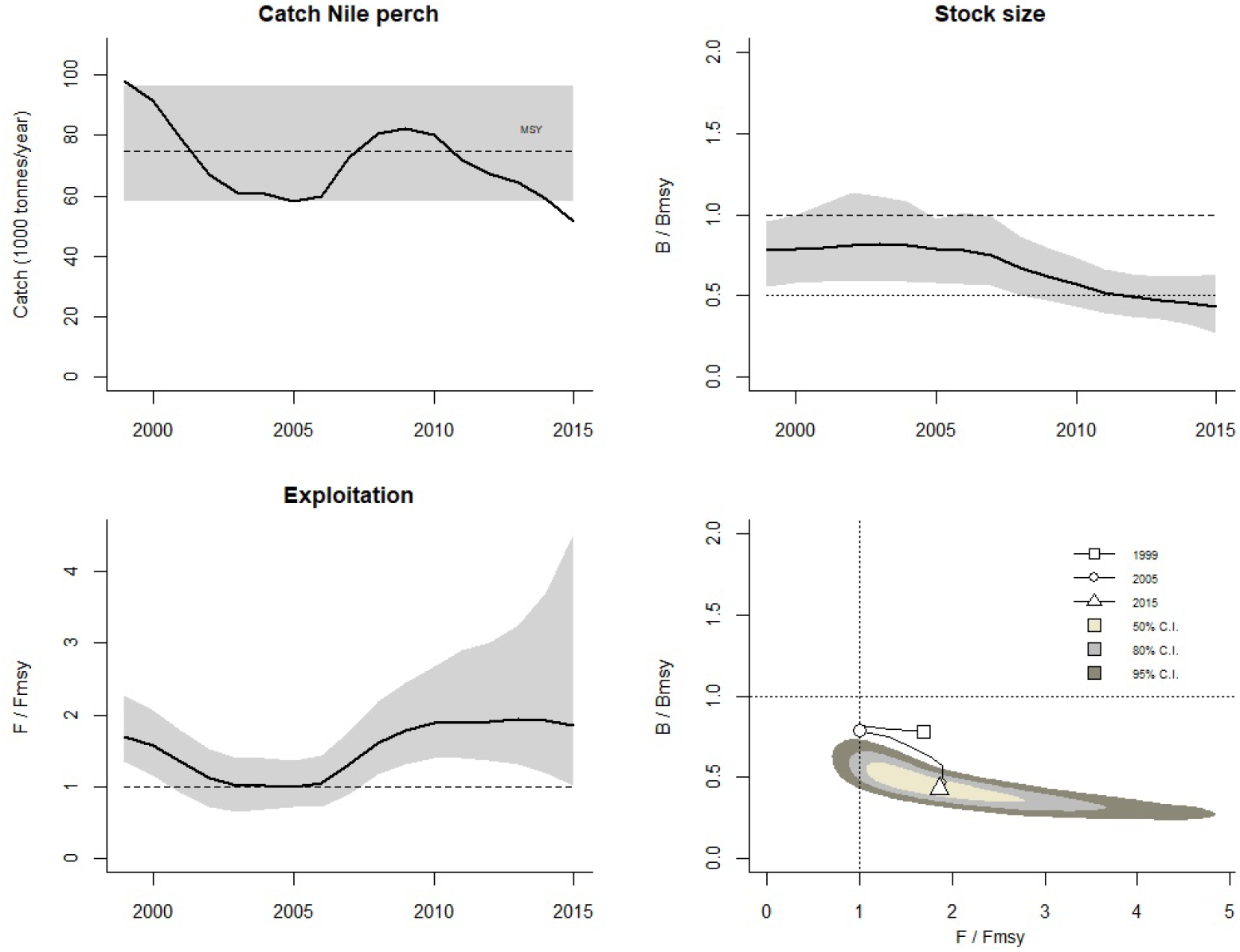
Trends in key management aspects of the Nile perch fishery in Lake Victoria, Uganda. The graphs show catches relative to MSY, with 95% confidence limits in grey (upper left); predicted relative total biomass (B/B_msy_) with the grey area indicating uncertainty (upper right); relative exploitation (F/F_msy_) and corresponding 95% confidence limits in grey (lower left); and stock status in relation to B/B_msy_ as a function of F/F_msy_ for the first (1965), intermediate (1989) and final (2015) years of assessment. The 50, 80 and 95% are Confidence levels around the assessment of the final year.

**Figure 3.**
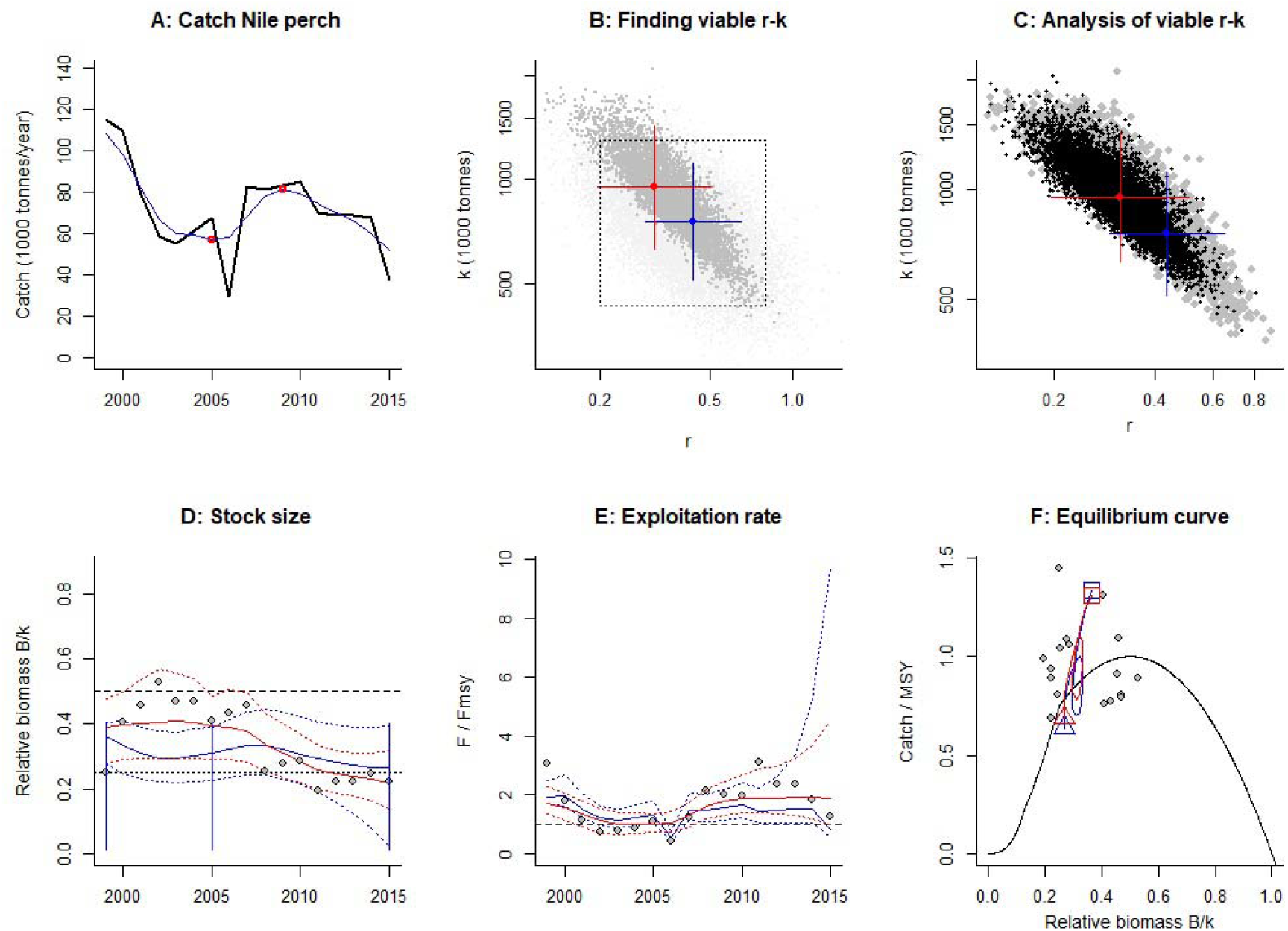
Results of the Nile perch fishery in Lake Victoria, Uganda. A shows time series in catch (black curve) and smoothed data with indication of highest and lowest catch (red curve). In B to F, red refers to estimates of BSM and blue to estimates of CMSY+. The crosses in B show the best r-k estimate of either methods (point in the center) and their 95% confidence limits (horizontal and vertical error bars). In dark grey are the pairs found to be compatible with the catches and biomass. In C, the black and dark grey dots are the viable r-k pairs found by BSM and CMSY respectively, with indication of crosses for best estimates with 95% confidence limits. Curves in D show the BSM and CMSY+ predictions of biomass, the dots the biomass data scaled by BSM, the vertical blue lines the prior biomass ranges. E shows the predictions for exploitation and catch per biomass as scaled by BSM (dots). The curves in F show the BSM and CMSY+ predictions of Schaufer equilibrium curves for catch/MSY relative to stock size (B/k) from the first (square) to the last year (triangle) of assessment, with the dots showing predicted catch per predicted biomass as scaled by BSM.

**Figure 4.**
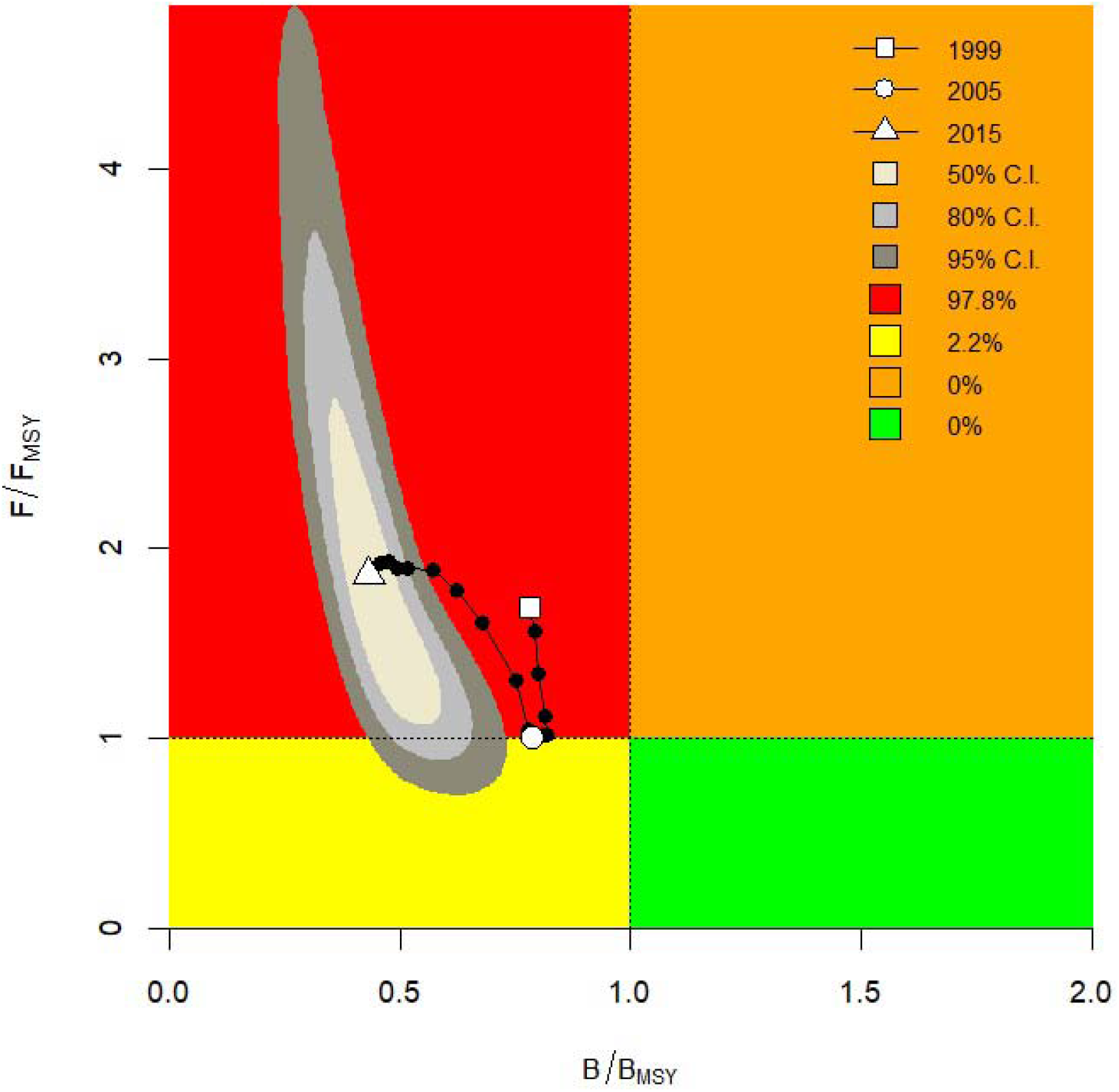
A Kobe plot for the Nile perch in the Lake Victoria, Uganda based on CMSY+ estimates of B/B_msy_ and F/F_msy_. A stock in the orange area is health but vulnerable to depletion by overfishing. In the red area, a stock is overfished and is undergoing overfishing, with too low biomass levels to produce maximum sustainable yields (MSY). In the yellow area, a stock is under reduced fishing pressure but recovering from too low biomass levels. The green area is the target area for management, indicating sustainable fishing pressure and healthy stock size capable of producing high yields close to MSY. The probabilities of the Nile perch stock being in any of these areas are given. the last year falling into one of the colored areas. The 50, 80 and 95% are confidence levels around the year of final assessment. The legend in the upper right graph also indicates.

**Table 2.** Lake wide estimates when abundance is fishery independent CPUE. Estimates are on based BSM with approximate 95% confidence limits in parentheses. Estimates for F_msy_, MSY and B_msy_ are long-term averages while others are for the last year in the dataset (2015).

| Stock | $F_{msy}$ (1/year) | MSY (1000 tonnes/year) | $B_{msy}$ (1000 tonnes) | B (1000 tonnes) | F (1/year) | Exploitation ($F/F_{msy}$) |
| --- | --- | --- | --- | --- | --- | --- |
| Nile tilapia | 0.0514(0.0308-0.0858) | 72.8(52 - 102) | 446(297-668) | 70.1(36.6-170) | 0.531(0.219-1.02) | 10.3(1.79-39.9) |
| Nile perch | 0.228(0.135-0.384) | 246(206-294) | 1080(675-1727) | 547(365-774) | 0.37(0.262-0.555) | 1.66(1.09-3.65) |
| Dagaa | 0.474(0.344-0.652) | 461(396 - 536) | 973(700-1351) | 1112(809-1361) | 0.484(0.395-0.665) | 1.03(0.781-1.46) |

In Uganda, the stock status of Nile tilapia and Nile perch was recruitment impaired, and Dagaa health. Nile tilapia stock was tending more towards collapsed status while Nile perch and Dagaa stocks were tending more towards overfished status (Table 4). For the whole lake, the stock status of Nile tilapia, Nile perch and Dagaa was collapsed, overfished and health respectively. The Nile perch and Dagaa stocks were respectively tending more towards recruitment impaired and overfished status (Table 4).

Our results confirm widespread overfishing for Nile tilapia and Nile perch. The poor status corresponds to poor fishing practices and intensive fishing pressure that have characterized the fisheries of the stocks for a longtime (Nyamweya et al. (2020). Njiru et al. (2007) assessed the two stocks in Kenya and observed high recruitment overfishing with 98% of Nile perch and 60% of Nile tilapia landed immature, high fishing mortality rate and degradation in life history. The degradation in life history in response to intensive fishing was found to be lake wide (Njiru et al. 2008).

The high fishing pressure and its persistence in the lake are consistent with our estimates of exploitation. In Uganda, exploitation has been above the reference level since 2006 for Nile perch (Figure 2 bottom right; Figure 3E) and 2003 for Nile tilapia (supplementary Figure 1 bottom right; supplementary Figure 2E). At around the same time, exploitation has since been above reference level for the stocks in the whole lake. As a result, we observed degradation in stock size depicted in declining B/B_msy_ (Figure 2 top right) and B/k (Figure 3D). See corresponding supplementary figures for other stocks.

Exploitation was higher for Nile tilapia stocks hence its worse status compared to Nile perch (Tables 2, 3 & 4). We presume that the Nile perch stock status would be worse than it is were it not for its high fecundity and pseudo protected areas offshore where fishing is probably restricted by distance and severity of weather. Nile perch individuals can obtain absolute fecundity of 16.8 million depending on size (Ogutu-ohwayo, 1988). This contrasts with the Nile tilapia whose mean absolute fecundity is 837 (Natugonza et al. 2016). The poor status of the two stocks is collectively illustrated in the Kobe plots which indicate that the stocks are 97.8% (Nile perch) and 100% (Nile tilapia) unsustainable in Uganda (Figure 4; supplementary figure 3) and 99.9% (Nile tilapia) and 99.1% (Nile perch) in the whole lake, requiring urgent management interventions

Dagaa, unlike the other stocks had health stock status in Uganda and the whole lake. The health status is depicted in low exploitation which has mainly been below the reference level (supplementary Figure 4 bottom right; supplementary Figure 5E; supplementary Figure 13 bottom right; supplementary Figure 14E). The health stock status cannot be attributed to good management which has been limited in the lake (Njiru et al. 2007). Indeed, fishing pressure on the stock has intensified because the number and panels of seines used to target it have increased while mesh sizes have declined (Mangeni-Sande et al. 2019). We attribute the better status of the stock to the high resilience of the species (Froese & Pauly, 2019). Dagaa has ability to double its biomass in <15 months and this is likely the reason it can bounce back from the high fishing pressure. Nevertheless, management needs to pay attention because the fishery is 17.7-30.2% unsustainable (supplementary figures 6 & 15) and tending more towards overfished status in both the whole lake and Ugandan part (Table 4).

Our estimates of MSY provide with managers, the fisheries potential of the stocks under good management. The MSY estimates for Nile perch and Nile tilapia were more than the most recent catches in the Ugandan part and the whole lake (Table 5). In Uganda, rebuilding the Nile tilapia and Nile perch stocks to MSY level could increase the catches of the stocks by 9.2% and 29.5% above the most recent catches respectively. At the whole lake level, the same intervention could increase the catches of the two stocks by 72.8% and 15.1% respectively (Table 5). To realize these benefits, managers should, in management objectives include rebuilding biomass of Nile tilapia and Nile perch to B_msy_ levels which were in all cases more than the current (2015) biomass (Table 2, 3 & 4). Estimates of biomass available for Nile perch since 2017 when enforcement was strengthened show that this is possible through eliminating illegal fishing practices (Hydro-acoustics Regional Working Group, 2019).

**Table 3.** Estimates for the Ugandan part of the lake. Estimates are based on BSM with approximate 95% confidence limits in parentheses. Estimates for F_msy_, MSY and B_msy_ are long-term averages while others are for the last year in the dataset (2015).

| Stock | $F_{msy}$ (1/year) | MSY (1000 tonnes/year) | $B_{msy}$ (1000 tonnes) | B (1000 tonnes) | F (1/year) | Exploitation ($F/F_{msy}$) |
| --- | --- | --- | --- | --- | --- | --- |
| Nile tilapia | 0.082(0.0501-0.134) | 19.9((15.7-25.4) | 121(80.5-183) | 30.4(16.3-53.4) | 0.556(0.316-1.03) | 6.77(2.09-24.4) |
| Nile perch | 0.136(0.0846-0.219) | 74.8(58-96.4) | 476(316-716) | 206(130-302) | 0.25(0.171-0.396) | 1.86(1.01-4.52) |
| Dagaa | 0.433(0.312-0.6) | 95.4(80.6-113) | 220(158-306) | 242(164-310) | 0.446(0.348-0.656) | 1.04(0.738-1.62) |

**Table 4.**
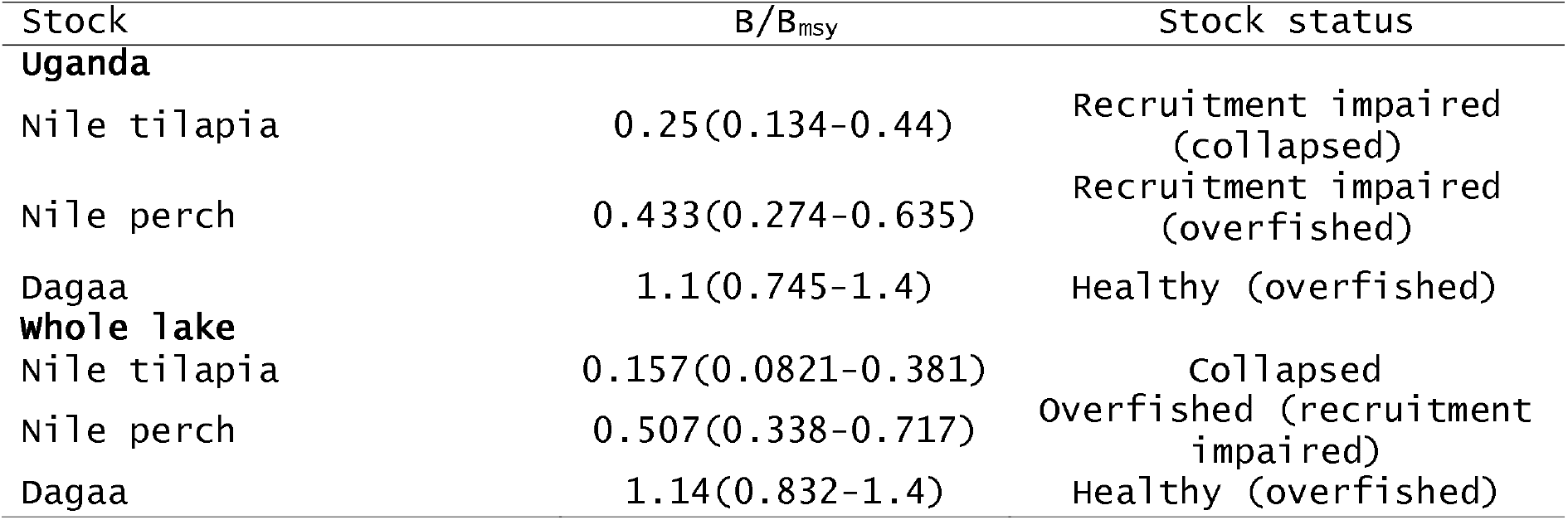
Stock status based on the classification of *B/B_msy_* values by Palomares et al. 2018. In parentheses is the stock status each of the stocks is tending more towards.

| Stock | $B/B_{msy}$ | Stock status |
| --- | --- | --- |
| **Uganda** |  |  |
| Nile tilapia | 0.25(0.134-0.44) | Recruitment impaired (collapsed) |
| Nile perch | 0.433(0.274-0.635) | Recruitment impaired (overfished) |
| Dagaa | 1.1(0.745-1.4) | Healthy (overfished) |
| **whole lake** |  |  |
| Nile tilapia | 0.157(0.0821-0.381) | Collapsed |
| Nile perch | 0.507(0.338-0.717) | Overfished (recruitment impaired) |
| Dagaa | 1.14(0.832-1.4) | Healthy (overfished) |

**Table 5.**
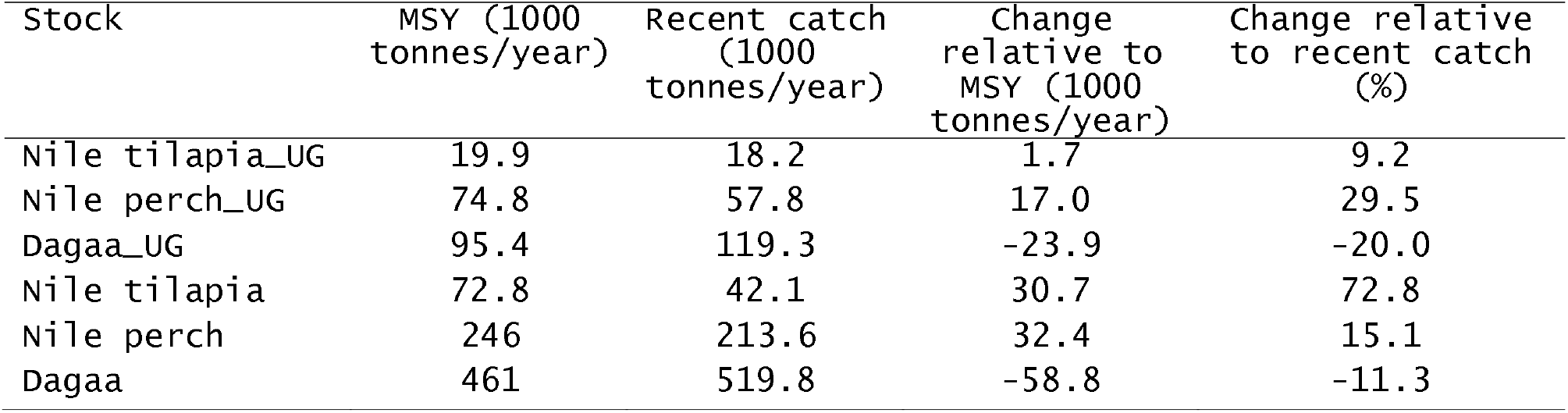
Estimates of MSY for the stocks in relation to recent catches (average for last three years (2013-2015).

The mean biomass of Nile perch in the whole lake in 2019 was 816,694 tonnes, with 705,458 tonnes as the lower limit and 940,922 tonnes as the upper limit. This was more than the biomass in 2015 and 24.4% less than B_msy_ (Table 3). In Uganda, the mean biomass was 422,076 tonnes with lower and upper limits of 366,757and 485,694 tonnes respectively which was also more than the 2015 biomass and only 11.3% less than B_msy_ (Table 2). These values indicate that the current biomass B_msy_ gap is closing faster in Uganda compared to the whole lake. The recovery for the Whole lake is probably constrained by Kenya which unlike Uganda and Tanzania, has not strengthened enforcement. Kenya should copy.

Unlike Nile tilapia and Nile perch, the MSY estimates for Dagaa were lower than recent catch. This means that much more is being taken than is supported by standing biomass, the same process that gradually brought about the observed poor status of the Nile tilapia and Nile perch stocks.

## Conclusion

The major objective of management on Lake Victoria should be rebuilding biomass of Nile tilapia and Nile perch to a level that can support catches at MSY. Eliminating illegal fishing practices has proved to be an effective way to achieve this because of the observed increase in biomass of Nile perch since 2017 when enforcement in Uganda and Tanzania was strengthened (Hydro-acoustics Regional Working Group, 2019). Kenya should do the same while Tanzania and Uganda should strengthen to close the gap between the current biomass and B_msy_.

After elimination of illegal fishing practices, the next task of management should be to ensure that catches remain low until biomass is ≥B_msy_ for at least three consecutive years (Froese et al. 2017).

After rebuilding, catches could be increased to MSY although a precautionary measure according to FAO is to exploit at the lower boundary of MSY (Tables 2 & 3) to guard against inefficiencies in enforcement and natural dynamics of fish stocks (Caddy et al. 1984). The precautionary measure could also cater for uncertainties in data due to unreported catches and cross border fishing and fish trade which are common on Lake Victoria (Heck et al. 2004). Cross border fishing and trade could lead to uncertainties in estimates of reference points especially at country level. For example, Kenyan fishers and traders who confessed to extensive cross-border fishing and trade give Kenya more catches than expected, a source of uncertainty (Geheb, 1997; Matsuishi et al. 2006). Indeed, our MSY estimates for Nile perch and Nile tilapia in Uganda (Table 3) are trumped by MSY estimates of 86,096 tonnes and 27,892 tonnes for the stocks respectively in Kenya (Aura et al. 2020). Cross border fishing and trade is the only plausible explanation for this.

To facilitate these interventions, routine data collection, preferably at an annual scale is indispensable to monitor biomass and catches to support stock assessments such as this to guide and evaluate management measures. Data collection has been a challenge particularly in Uganda. For instance, since 2015, no catch assessment surveys have been done in the Ugandan part of Lake Victoria which is regrettable.

## Supporting information

supplementary table and graphs

## Acknowledgement

We are grateful to funders who supported projects from which the data used in this study were generated.

## Notes

### Competing Interest Statement

The authors have declared no competing interest.

