## supplementary table and graphs for "Status and targets for rebuilding the three major fish stocks in Lake Victoria"

Table 1. Lake wide estimates when abundance is absolute biomass. Estimates are on based BSM with approximate 95% confidence limits in parentheses. Estimates for F_msy_, MSY and B_msy_ are long-term averages while others are for the last year in the dataset (2015).

| Stock | F_msy_ (1/year) | MSY (1000 tonnes/year) | B_msy_ (1000 tonnes) | B/B_msy_ | B (1000 tonnes) | F (1/year) | Exploitation (F/F_msy_) |
| --- | --- | --- | --- | --- | --- | --- | --- |
| Nile tilapia | 0.111(0.0665-0.184) | 71(59.1-85.3) | 357(237-539) | 0.278(0.123-0.465) | 99.3(43.9–166) | 0.399(0.238-0.902) | 3.57(1.27-19.3 |
| Nile perch | 0.201(0.106-0.38) | 219(171–281) | 1091(651–1827) | 0.599(0.219-0.929) | 653(239-1014) | 0.318(0.205-0.872) | 1.58(0.919-10.8) |
| Dagaa | 0.347(0.252-0.477) | 417(305–570) | 1202(914–1580) | 0.848(0.485 - 1.33) | 1019(583–1598) | 0.526(0.336-0.92) | 1.53(0.853-2.89) |

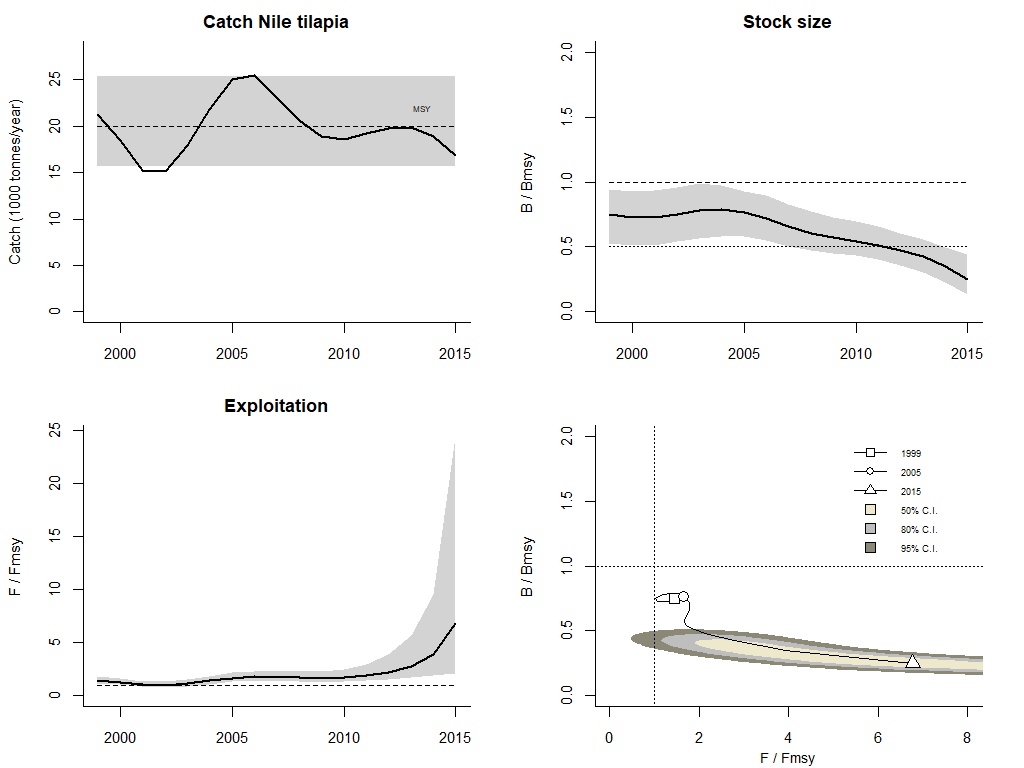

Figure 1 Trends in key management aspects of the Nile tilapia fishery in Lake Victoria, Uganda. The graphs show catches relative to MSY, with 95% confidence limits in grey (upper left); predicted relative total biomass (B/B_msy_) with the grey area indicating uncertainty (upper right); relative exploitation (F/F_msy_) and corresponding 95% confidence limits in grey (lower left); and stock status in relation to B/B_msy_ as a function of F/F_msy_ for the first (1965), intermediate (1989) and final (2015) years of assessment. The 50, 80 and 95% are Confidence levels around the assessment of the final year.

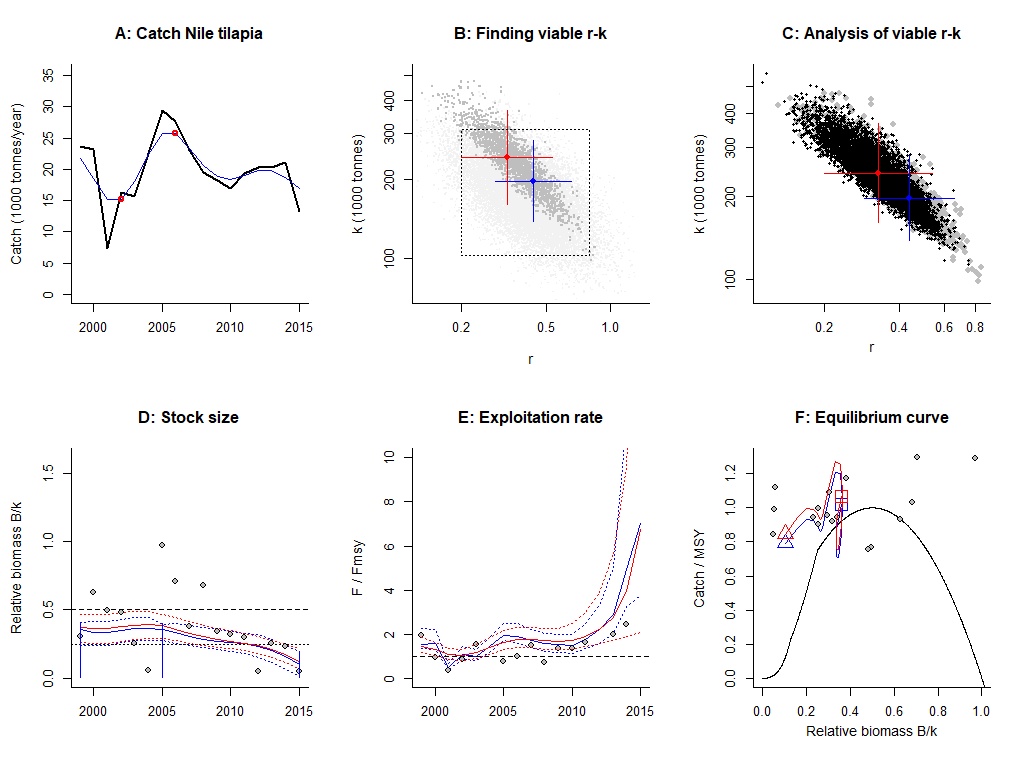

Figure 2 Results of the Nile tilapia fishery in Lake Victoria, Uganda. A shows time series in catch (black curve) and smoothed data with indication of highest and lowest catch (red curve). In B to F, red refers to estimates of BSM and blue to estimates of CMSY+. The crosses in B show the best r-k estimate of either methods (point in the center) and their 95% confidence limits (horizontal and vertical error bars). In dark grey are the pairs found to be compatible with the catches and biomass. In C, the black and dark grey dots are the viable r-k pairs found by BSM and CMSY respectively, with indication of crosses for best estimates with 95% confidence limits. Curves in D show the BSM and CMSY+ predictions of biomass, the dots the biomass data scaled by BSM, the vertical blue lines the prior biomass ranges. E shows the predictions for exploitation and catch per biomass as scaled by BSM (dots). The curves in F show the BSM and CMSY+ predictions of Schaufer equilibrium curves for catch/MSY relative to stock size (B/k) from the first (square) to the last year (triangle) of assessment, with the dots showing predicted catch per predicted biomass as scaled by BSM.

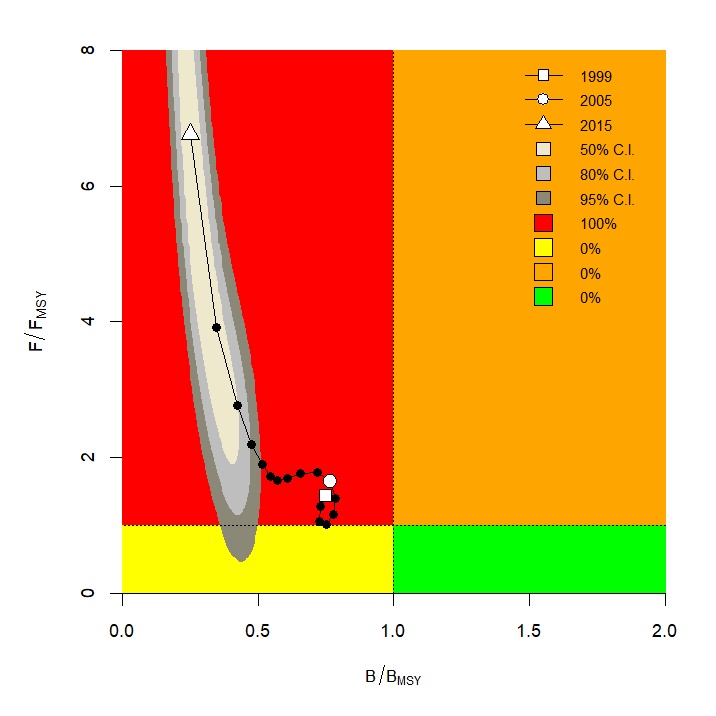

Figure 3 A Kobe plot for the Nile tilapia fishery in the Lake Victoria, Uganda based on CMSY+ estimates of B/Bmsy and F/Fmsy. A stock in the orange area is health but vulnerable to depletion by overfishing. In the red area, a stock is overfished and is undergoing overfishing, with too low biomass levels to produce maximum sustainable yields (MSY). In the yellow area, a stock is under reduced fishing pressure but recovering from too low biomass levels. The green area is the target area for management, indicating sustainable fishing pressure and healthy stock size capable of producing high yields close to MSY. The probabilities of the Nile tilapia stock being in any of these areas are given. the last year falling into one of the colored areas. The 50, 80 and 95% are confidence levels around the year of final assessment. The legend in the upper right graph also indicates.

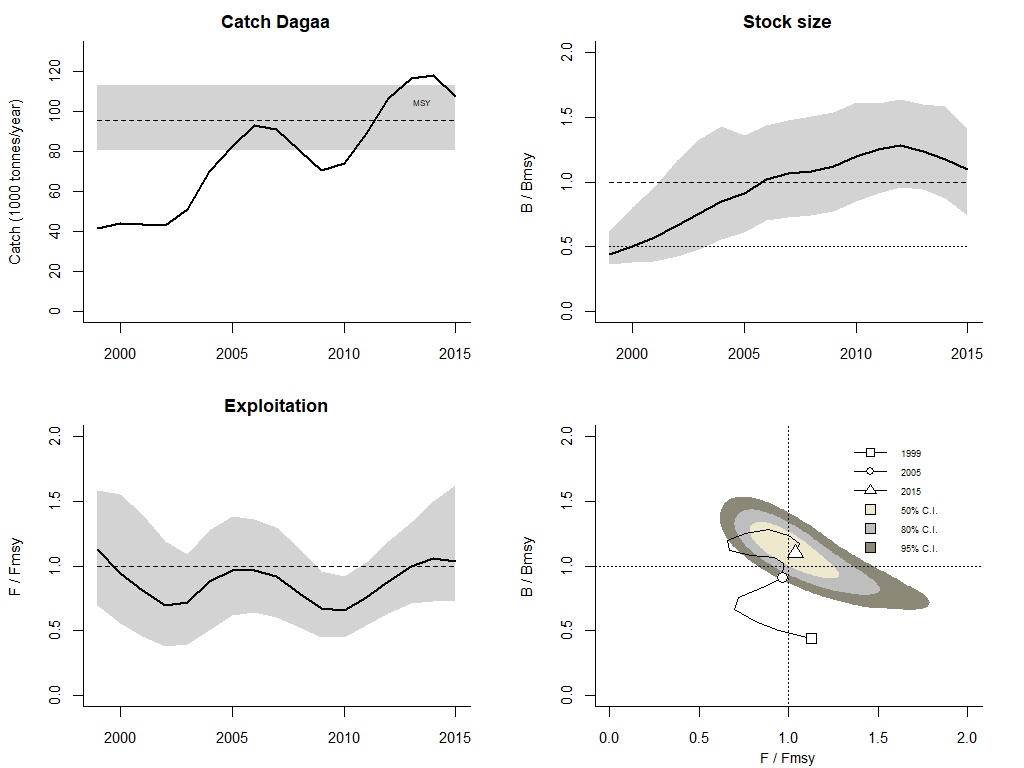

Figure 4 Trends in key management aspects of the Dagaa fishery in Lake Victoria, Uganda. Graph details are as in Figure 1

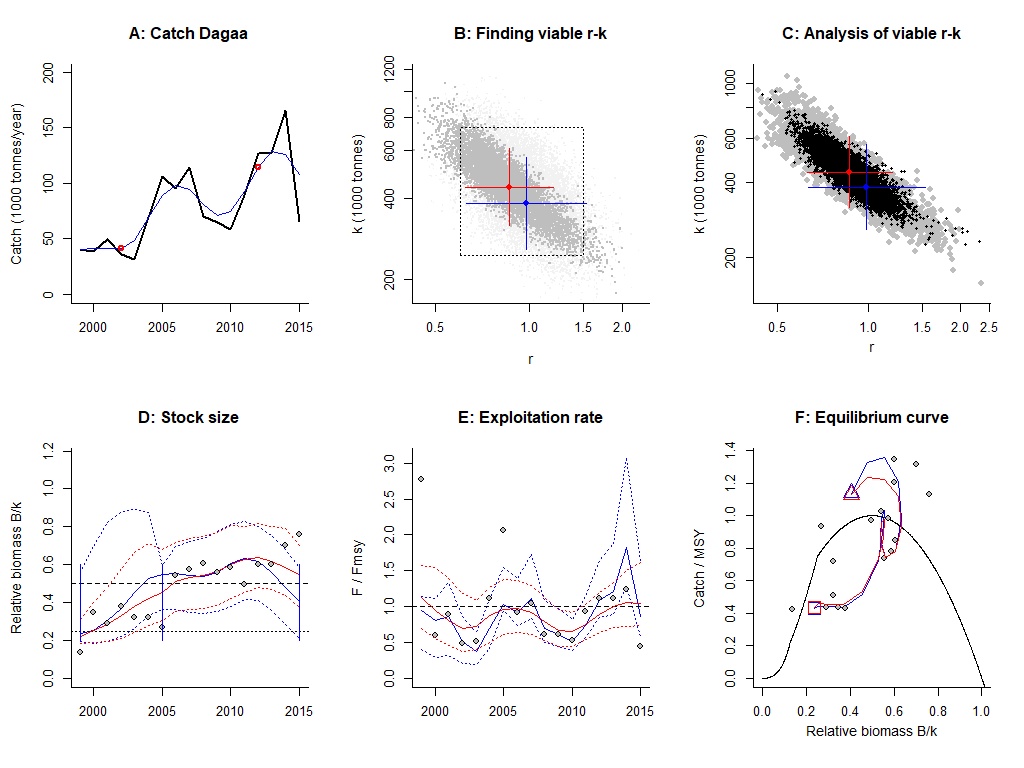

Figure 5 Results of the Dagaa fishery in Lake Victoria, Uganda. Details of the graphs are as in Figure 2

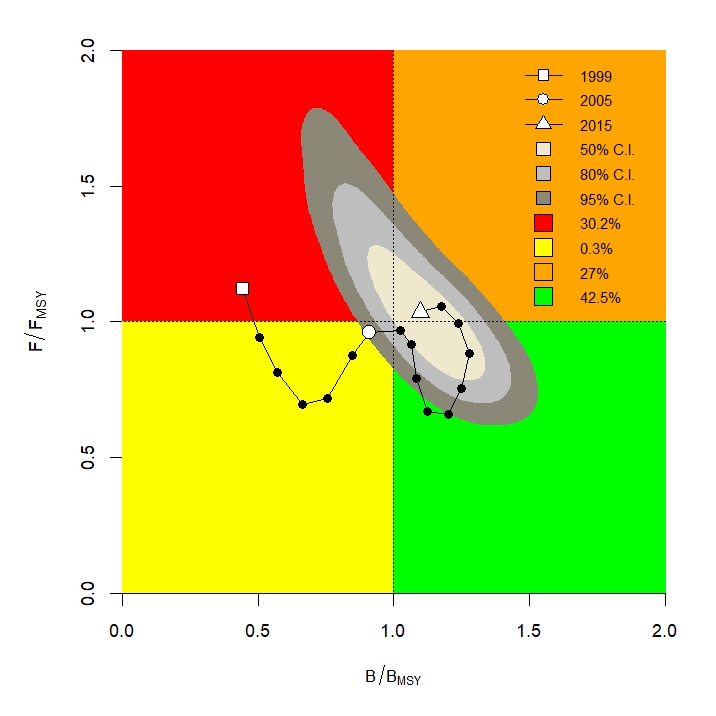

Figure 6 A Kobe plot for the Dagaa fishery in the Lake Victoria, Uganda based on CMSY+ estimates of B/Bmsy and F/Fmsy. Details are as in Figure 3

### **Whole lake when abundance is fishery independent CPUE**

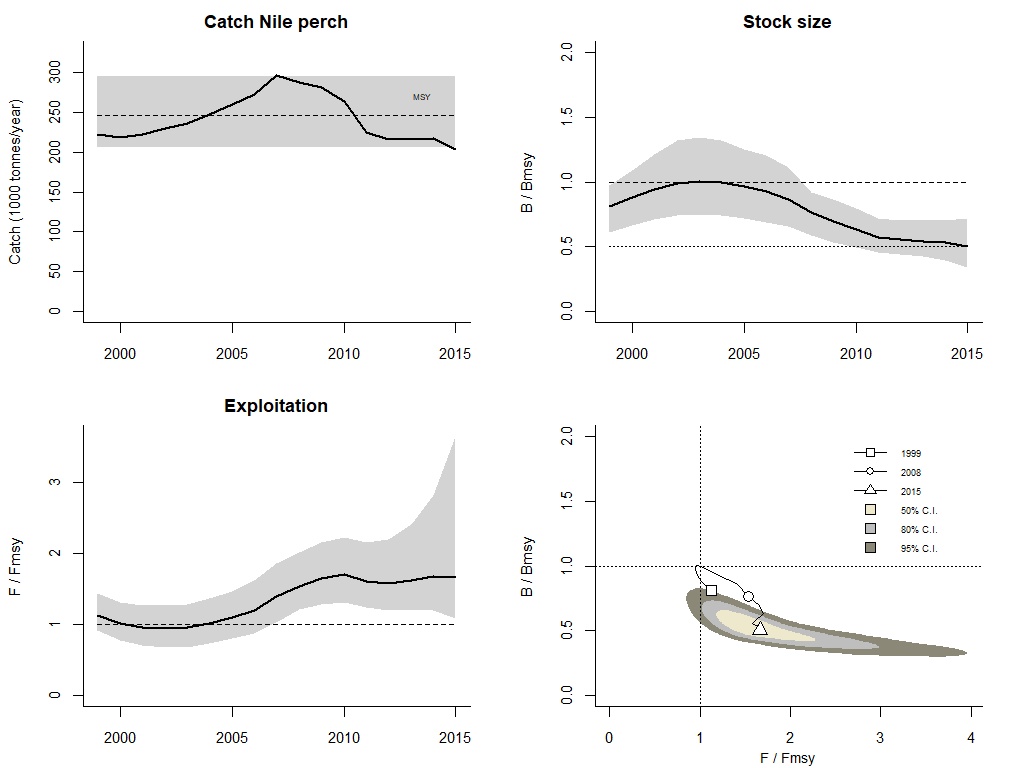

Figure 7 Trends in key management aspects of the Nile perch fishery in the whole of Lake Victoria. Graph details are as in Figure 1

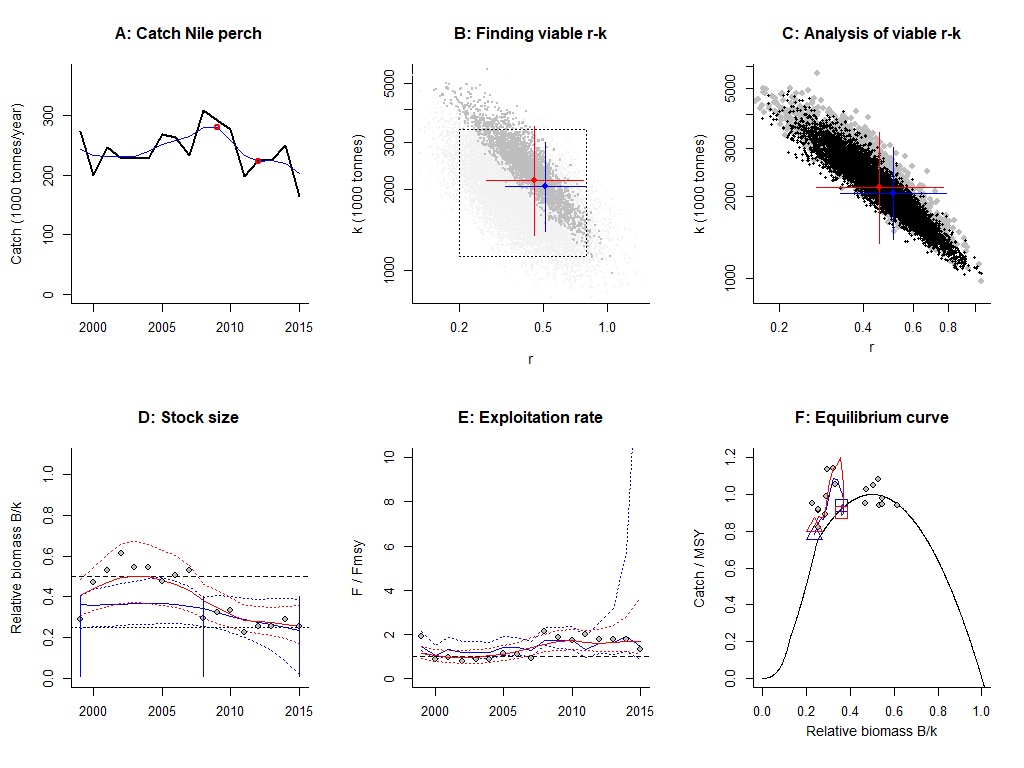

Figure 8 Results of the Nile perch fishery in the whole of Lake Victoria. Details of the graphs are as in Figure 2

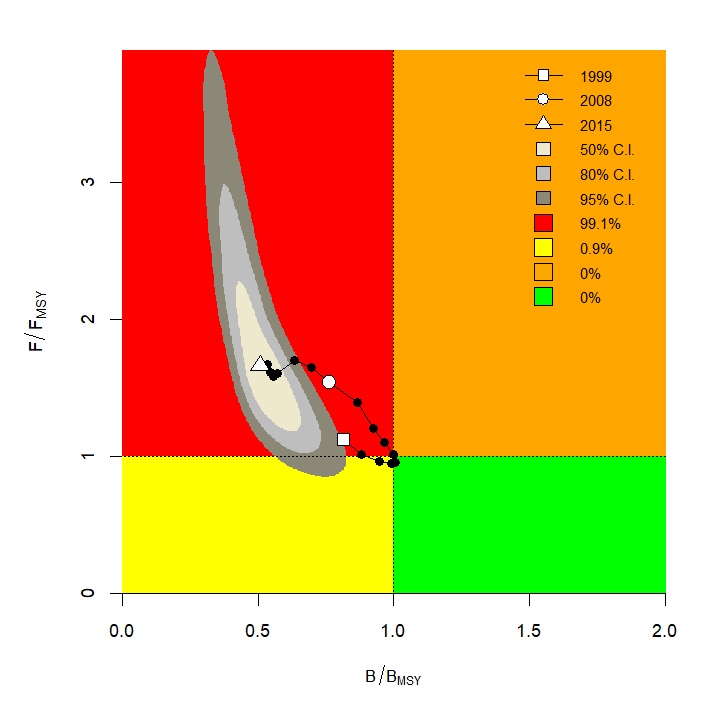

Figure 9 A Kobe plot for the Nile perch fishery in the whole of Lake Victoria based on CMSY+ estimates of B/Bmsy and F/Fmsy. Details are as in Figure 3

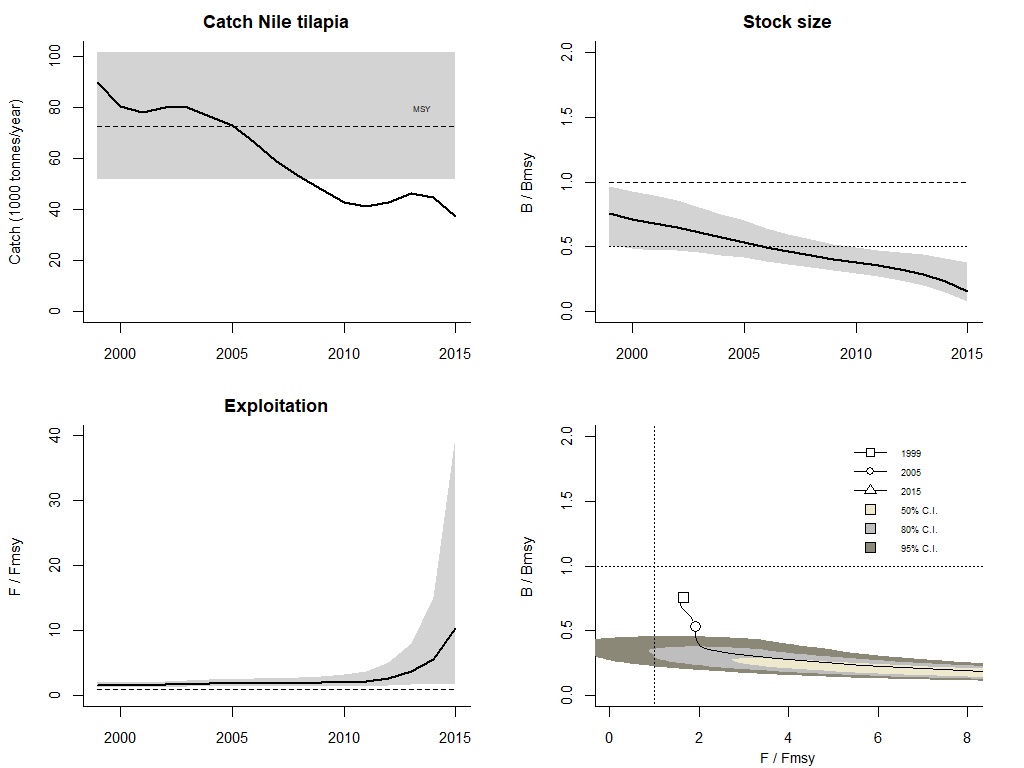

Figure 10 Trends in key management aspects of the Nile tilapia fishery in the whole of Lake Victoria. Graph details are as in Figure 1

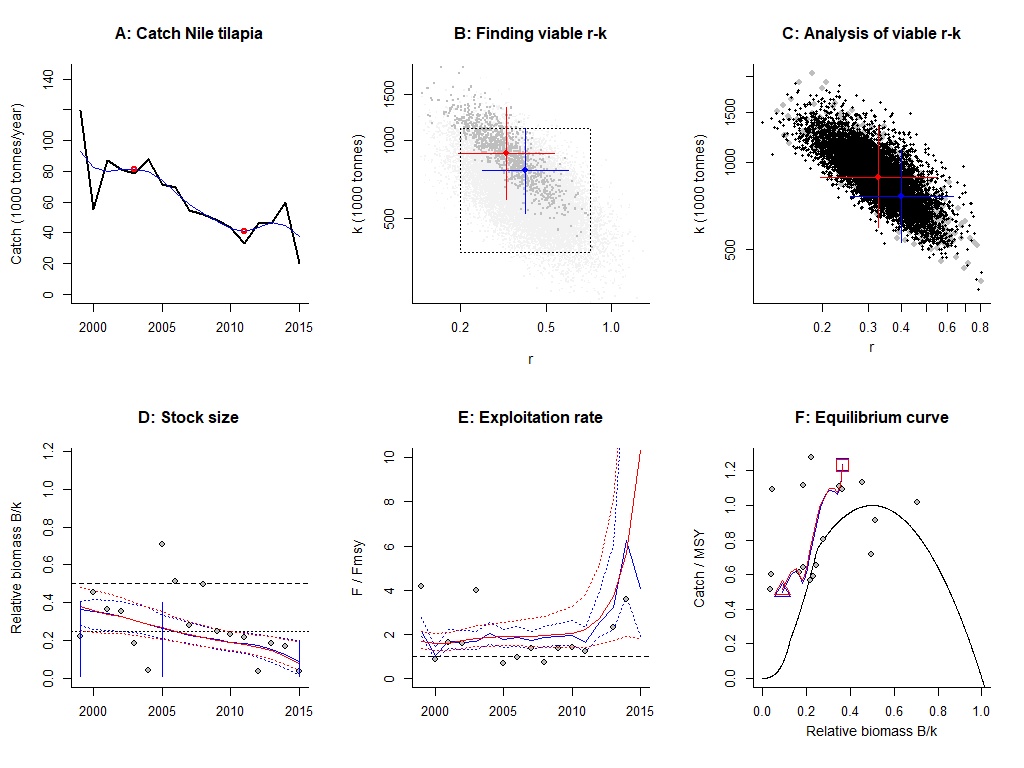

Figure 11 Results of the Nile tilapia fishery in the whole of Lake Victoria. Details of the graphs are as in Figure 2

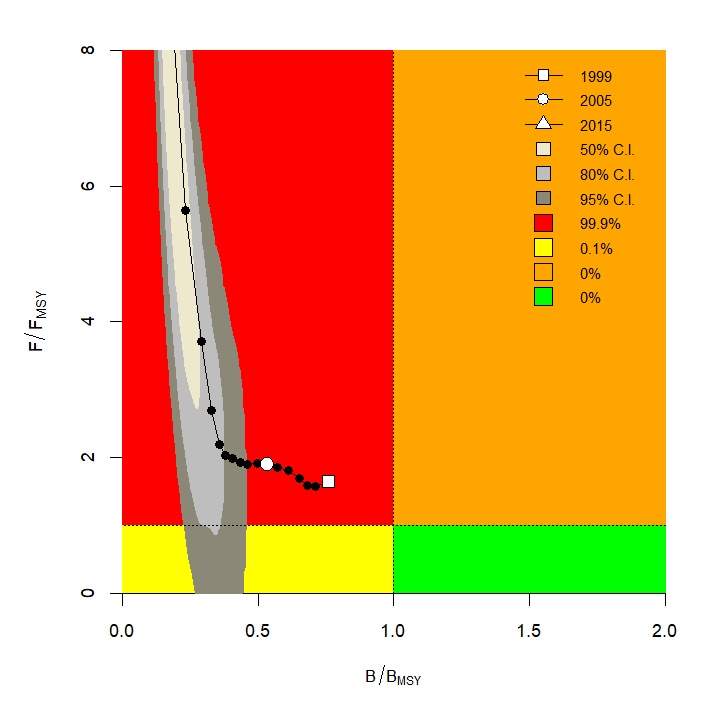

Figure 12 A Kobe plot for the Nile tilapia fishery in the whole of Lake Victoria based on CMSY+ estimates of B/Bmsy and F/Fmsy. Details are as in Figure 3

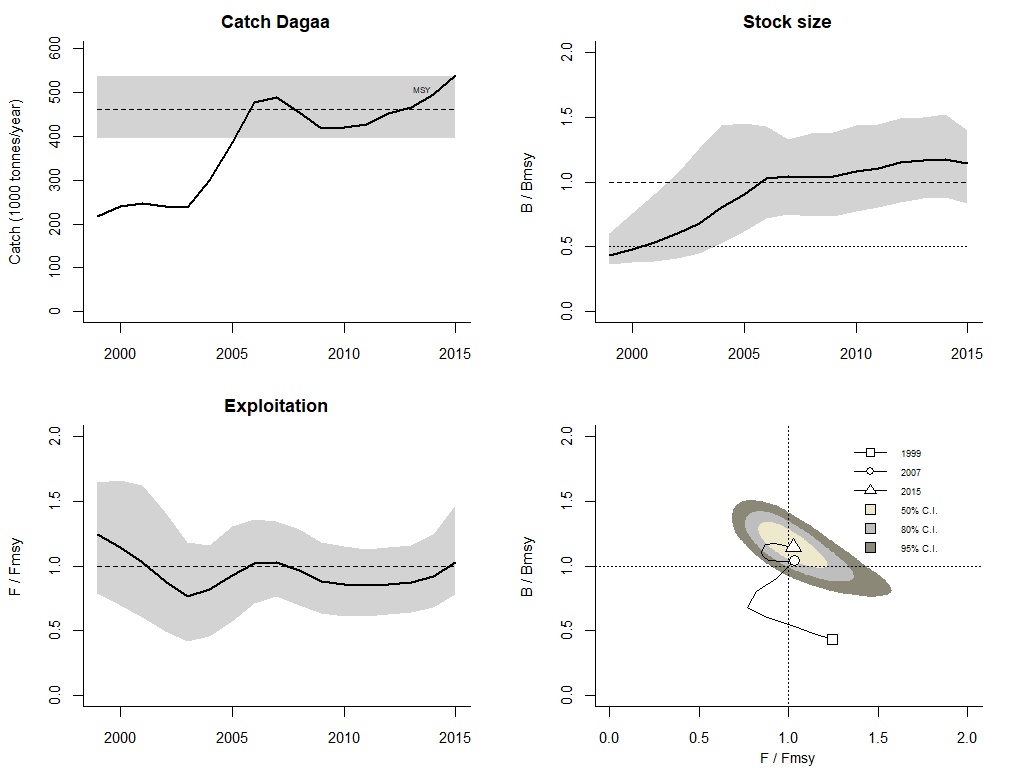

Figure 13 Trends in key management aspects of the Dagaa fishery in the whole of Lake Victoria. Graph details are as in Figure 1

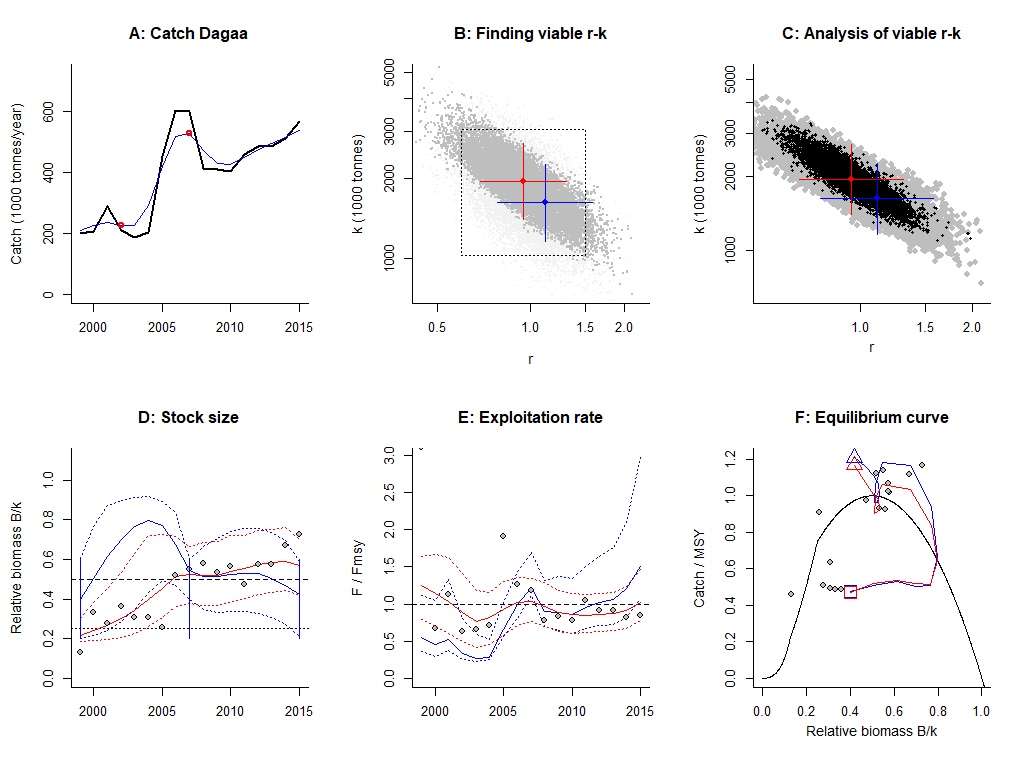

Figure 14 Results of the Dagaa fishery in the whole of Lake Victoria. Details of the graphs are as in Figure 2

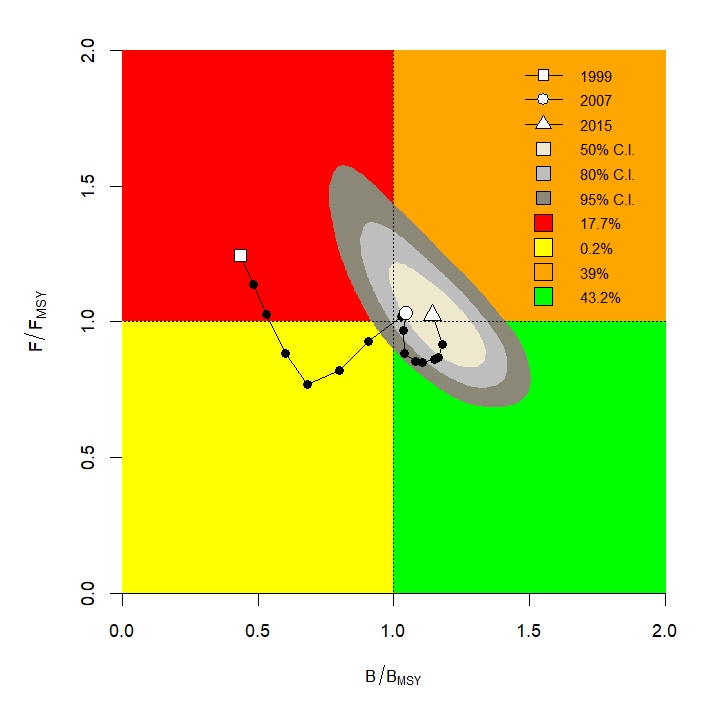

Figure 15 A Kobe plot for the Dagaa fishery in the whole of Lake Victoria based on CMSY+ estimates of B/Bmsy and F/Fmsy. Details are as in Figure 3
